# Plasma redox imbalance caused by albumin oxidation promotes lung-predominant NETosis and pulmonary cancer metastasis

**DOI:** 10.1101/273037

**Authors:** Minoru Inoue, Masahiro Enomoto, Yuhki Koike, Marco A. Di Grappa, Xiao Zhao, Kenneth Yip, Shao Hui Huang, John N. Waldron, Mitsuhiko Ikura, Fei-Fei Liu, Scott V. Bratman

## Abstract

Neutrophil extracellular traps (NETs) entrap circulating tumor cells (CTCs) and promote metastasis within distant organs in preclinical models^1,2^. In these models, NETosis is triggered by exogenous massive inflammatory stimuli, and thus it remains unknown whether cancer hosts under physiologic inflammation-free conditions experience NETosis and consequent cancer metastasis. Here we show that plasma redox imbalance caused by albumin oxidation promotes inflammation-independent NETosis and cancer metastasis specifically in the lungs. Albumin is the major source of free thiol that maintains redox balance *in vitro* and *in vivo*. Oxidation of albumin-derived free thiol is sufficient to trigger NETosis *via* accumulation of reactive oxygen species within neutrophils. The resultant NETs are found predominantly within lungs where they contribute to the colonization of CTCs leading to pulmonary metastases in mouse models. These effects are abrogated by pharmacologic inhibition of NET formation. Moreover, albumin oxidation and the resultant decline of plasma free thiol are associated with pulmonary metastasis in a cohort of head and neck cancer patients. These results implicate plasma redox balance as an endogenous and physiologic regulator of NETosis and pulmonary cancer metastasis, providing new therapeutic and diagnostic opportunities for combatting cancer progression.

## Introduction

Neutrophils release web-like DNA-containing structures (i.e., neutrophil extracellular traps [NETs], generated by a process termed NETosis) to entrap pathogens and prevent their dissemination into the circulation^3^. There is emerging evidence from preclinical cancer models that this entrapment effect of NETs is a pathological mechanism for metastasis initiation by promoting colonization of circulating tumor cells (CTCs)^1,2^. These models involve massive systemic inflammation, for example by injection of lipopolysaccharide (LPS)^4^, clamping of hepatic artery and portal vein^2,5^, or cecum ligation^1^. As these perturbations do not faithfully reproduce the clinical state of cancer patients and are non-specific in their mechanisms of inducing NETosis, it remains unknown whether NETs have a major influence on distant metastasis in human cancer. Thus, in the absence of massive systemic inflammation, we sought to identify endogenous and physiologic regulators of NETosis and their role in cancer metastasis.

Accumulation of the reactive oxygen species (ROS) within neutrophils is a key process for the initiation of NETosis^6^. Inhibiting this process by antioxidants such as free thiol-containing agents prevents NETosis^7-9^. Amongst the endogenous antioxidants, albumin comprises the largest plasma thiol pool^10^ and thus determines the plasma redox status. The interaction of the free thiol group in albumin with ROS produces oxidized albumin in exchange for catabolizing ROS^11^. Accumulation of oxidized albumin is associated with the progression in various chronic diseases^12-14^, but there is no known role for albumin oxidation in the pathophysiology of cancer. Here, we investigated the impact of plasma redox balance maintained by albumin thiol on NETosis and cancer metastasis.

## Results

### Albumin oxidation causes ROS accumulation within neutrophils and NETosis without inflammation-related stimuli

In order to investigate a possible role of albumin in regulating NETosis *in vitro*, we examined the effects of albumin depletion on cultured neutrophils. Human neutrophils were cultured under albumin-containing and albumin-free conditions using medium supplemented with 1% fetal bovine serum (FBS; 1% FBS), no FBS (0% FBS), or 1% albumin-depleted FBS (1% FBS-Alb). The 0% FBS and 1% FBS-Alb conditions displayed approximately half of the free thiol levels compared to 1% FBS (Fig. 1a). Six-hour incubation under both albumin-free conditions led neutrophils to form NETs, characterized by elongated DNA fibers (Fig. 1b) in association with LL-37 (Extended Data Fig. 1), which is identical to NETs triggered by inflammation-related stimuli with phorbol 12-myristate 13-acetate (PMA)^15^. Consistent with this observation, levels of extracellular DNA released into the culture medium were elevated in both albumin-free conditions (Fig. 1c), and the extracellular DNA was physically associated with LL-37 (Fig. 1d, e). Further supporting the role of albumin in regulating NETosis, a free thiol-containing agent, *N*-acetylcysteine (NAC)^7,8^, and an inhibitor of peptidyl arginine deiminase (PAD), Cl-amidine^16,17^, both blocked NETosis under albumin-free conditions (Extended Data Fig. 2a-d); these results indicate that loss of albumin induces NETosis through the established mechanisms of intracellular ROS accumulation and PAD activation.

**Figure 1.**
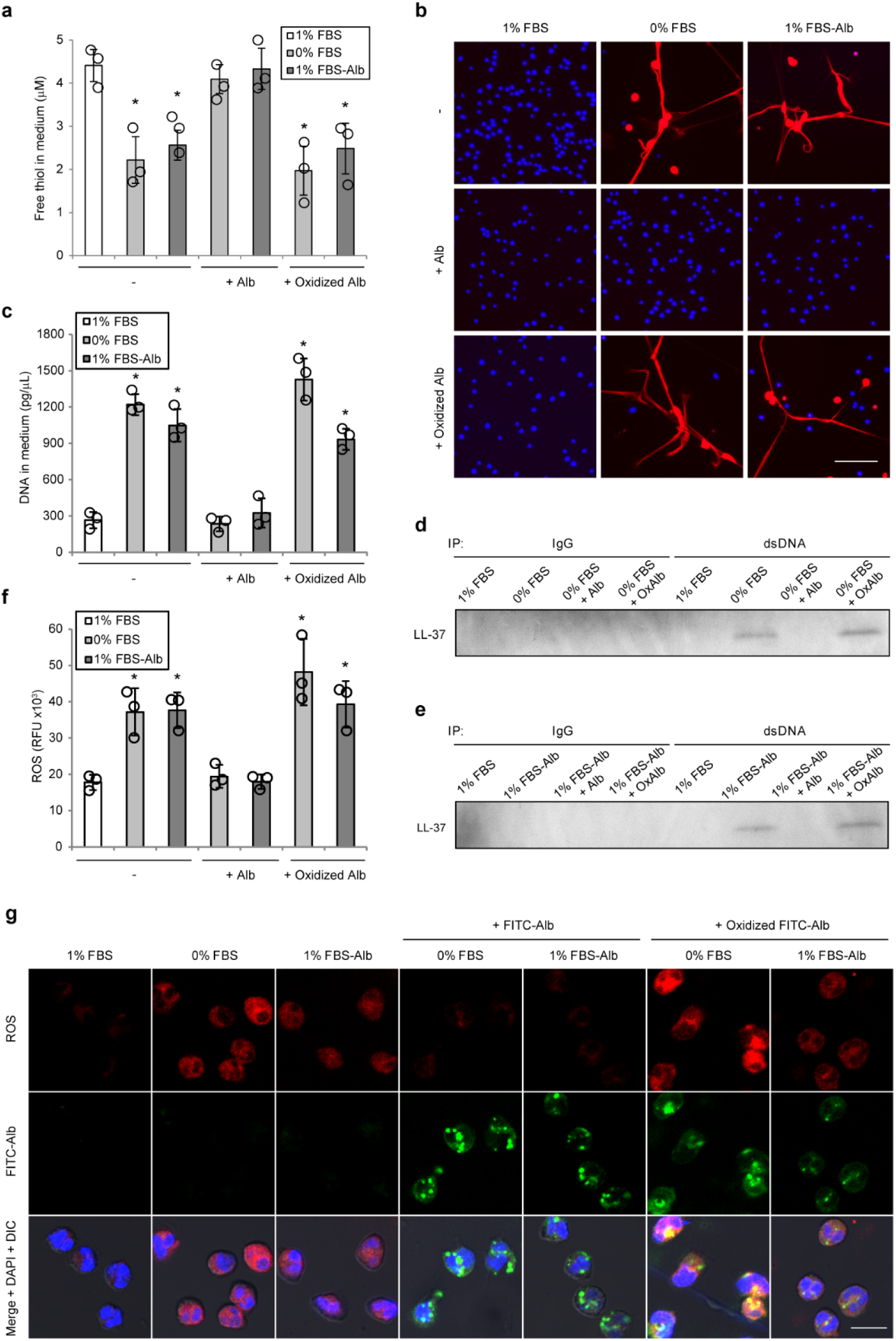
Albumin oxidation triggers NETosis *via* intracellular ROS accumulation. (a) Free thiol concentration in medium containing 1% FBS, 0% FBS, and 1% albumin-depleted FBS (1% FBS-Alb) with or without albumin replenishment (final concentration: 0.02 g/dL, matching the albumin concentration in the condition of 1% FBS). (b-g) Human neutrophils were cultured under the conditions of 1% FBS, 0% FBS, and 1% FBS-Alb with or without albumin replenishment for 6 hours (b-e), 5 minutes (f), or 1 hour (g). (b) Representative images of neutrophils stained with cell-permeable DNA dye, Hoechst 33342 (blue), and cell-impermeable DNA dye, SytoxOrange (red). Bar = 50 μm. (c) Concentration of extracellular DNA within culture medium. (d, e) Detection of LL-37 physically associated with extracellular DNA within culture medium. Extracellular DNA was immunoprecipitated from culture medium using an anti-dsDNA antibody or IgG isotype control, and the isolated DNA was subjected to Western blotting for LL-37. (f) Intracellular ROS levels within human neutrophils in the indicated culture conditions. (g) Representative images of intracellular ROS and albumin in neutrophils. Intracellular ROS was stained by CellROX Deep Red Reagent. Albumin internalization by neutrophils was visualized using albumin-fluorescein isothiocyanate conjugate (FITC-Alb). Bar = 10 μm. (a, c, f) Results represent individual values with the mean ± s.d. (*n* = 3; biological triplicates, \**P*<0.05 by Dunnett’s test).

Based on its known capabilities for ROS scavenging, we reasoned that the free thiol of albumin might be required for inhibiting NETosis. Indeed, although replenishment of bovine serum albumin (BSA) effectively inhibited NETosis under albumin-free conditions (Fig. 1b-e), BSA oxidized by chloramine-T^18^ (Fig. 1a) was incapable of inhibiting NETosis (Fig. 1b-e). Intracellular ROS level, which was elevated in neutrophils cultured under albumin-free conditions (Fig. 1f,g), was maintained at a comparably low level by BSA but not by oxidized BSA (Fig. 1f, g), although uptake of BSA and oxidized BSA by neutrophils was similarly observed (Fig. 1g). Taken together, free thiol supplied by albumin maintains redox balance in neutrophils and its oxidation triggers NETosis without inflammation-related stimuli.

### Plasma redox imbalance represented by albumin deficiency or oxidation triggers inflammation-independent NETosis

We next examined whether loss of albumin free thiol could be a non-inflammatory trigger for NETosis *in vivo*. First, we assessed the contribution of albumin to total plasma free thiol in mice. Compared with wild-type C57BL/6 mice, mice that were genetically deficient for albumin (Alb^−/−^)^19^ contained approximately 90% less total plasma free thiol (Fig. 2a). Albumin depletion from plasma obtained from other mouse strains (i.e., NOD scid gamma [NSG] and BALB/cByJ) confirmed that the vast majority of plasma free thiol was supplied by albumin (Fig. 2a). These results indicate that free thiol provided by circulating albumin determines plasma redox balance in mice.

**Figure 2.**
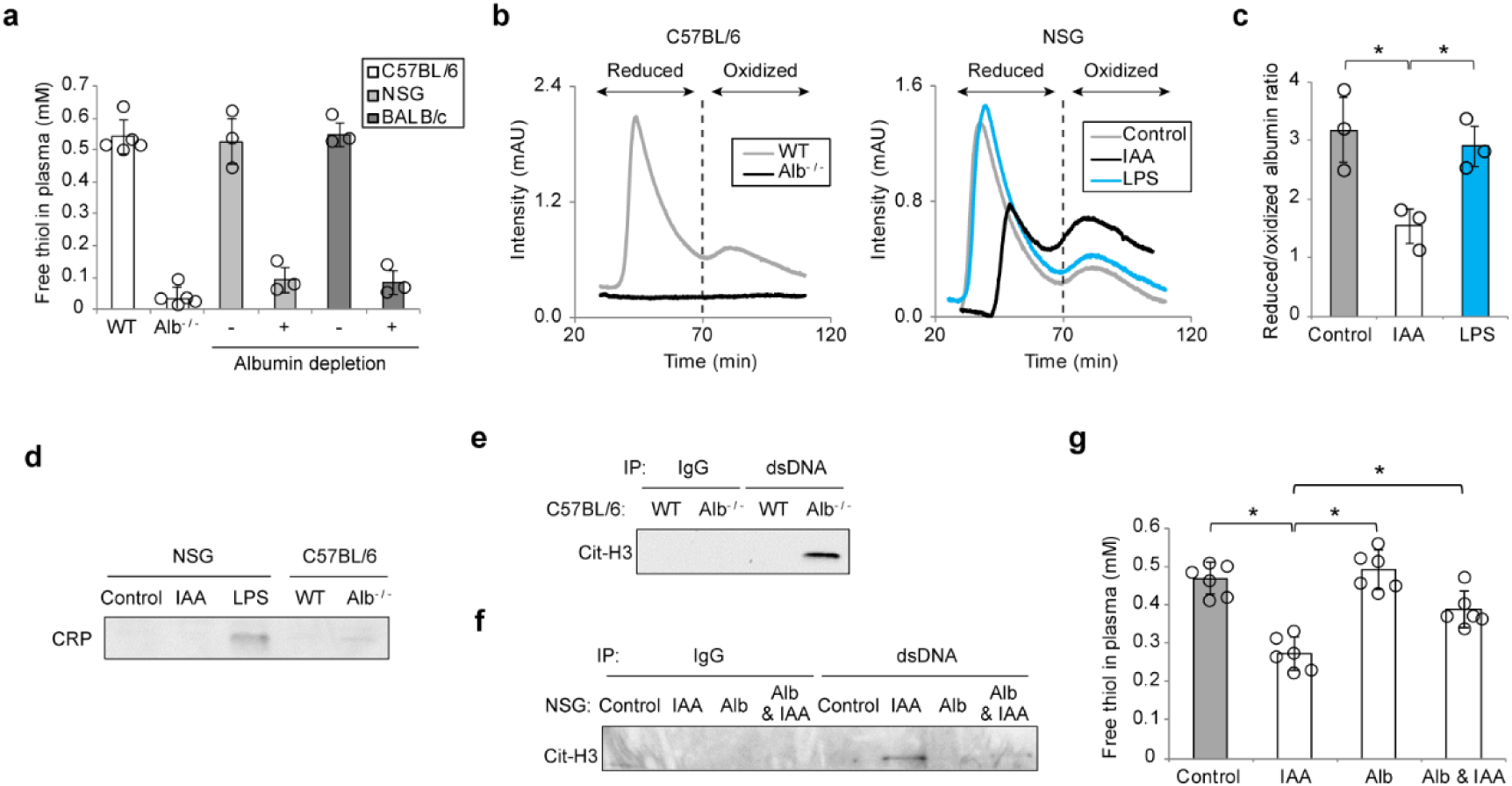
Plasma redox imbalance due to albumin deficiency or oxidation induces inflammation-independent NETosis. (a) Concentration of free thiol in plasma from wild-type (WT) vs. albumin deficient (Alb^−/−^) C57BL/6 mice and control vs. albumin-depleted plasma from NOD scid gamma (NSG) and BALB/c mice. (b-d) Plasma isolated from the blood in WT and Alb^−/−^ C57BL/6 mice and NSG mice 2 days post saline, IAA (30 mg/kg), or LPS (8 mg/kg) injection was subjected to FPLC (b, c) and Western blotting (d). (b) Representative FPLC results for the redox state of albumin in C57BL/6 (left) and NSG (right) mice. (c) The ratio of peak intensity for reduced/oxidized albumin in NSG mice. (d) Western blotting with an antibody against C-reactive protein (CRP). (e-g) Plasma was isolated from the blood in WT and Alb^−/−^ C57BL/6 mice and NSG mice 2 days post saline or IAA (30 mg/kg) injection following intravenous injection of PBS or murine albumin (20 mg/mouse). (e, f) Immunoprecipitation of cell-free DNA in the plasma from C57BL/6 (e) and NSG (f) mice using an anti-dsDNA antibody followed by Western blotting using an antibody against citrullinated histone H3 (Cit-H3). (g) Plasma free thiol concentration. (a, c, g) Results represent individual values with the mean ± s.d. (*n* = 3-6; biological replicates, \**P*<0.05 by Dunnett’s test).

Next, we tested the association between loss of albumin free thiol and NETosis. For this purpose, we used two different mouse models, Alb^−/−^ mice and Alb^wt/wt^ mice with pharmacological depletion of albumin free thiol by iodoacetamide (IAA)^20,21^. *In vitro*, IAA-treated BSA was incapable of scavenging ROS within neutrophils and inhibiting resultant NETosis (Extended Data Fig 3a-d), in keeping with the effects of albumin oxidation (Fig. 1c, d). *In vivo*, IAA administration reduced plasma free thiol levels with the lowest level observed at day 2 following injection (Extended Data Fig. 4a) without significantly impacting plasma albumin concentration (Extended Data Fig. 4b). To obtain direct evidence of albumin oxidation by IAA, we performed fast protein liquid chromatography (FPLC) to analyze the redox state of albumin. This assay shows albumin-specific signal (Fig. 2b, left) and provides its redox profile (Extended Data Fig. 5a-c). Compared to control mice, plasma obtained at day 2 from IAA-injected mice contained less reduced and more oxidized albumin (Fig. 2b, right; Fig. 2c). This oxidative shift of albumin was not observed in the plasma from mice in which systemic inflammation was induced by lipopolysaccharide (LPS) injection (Fig. 2b, right; Fig. 2c). As expected, LPS injection in these mice led to elevated plasma levels of C-reactive protein (CRP), a commonly used marker of inflammation; in contrast, neither genetic knockout of albumin nor pharmacological depletion of albumin free thiol led to elevated plasma levels of CRP (Fig. 2d). We further identified evidence of NETosis in the plasma from Alb^−/−^ mice and from IAA-injected mice by detecting a NET marker, citrullinated histone H3 (Cit-H3)^22^ in physical association with circulating cell-free DNA (Fig. 2e, f). When murine albumin was injected prior to IAA injection, IAA-evoked NETosis was suppressed along with normalization of plasma free thiol level (Fig. 2f, g), indicating that IAA triggers NETosis through albumin oxidation and consequent decrease in plasma free thiol. Collectively, plasma redox imbalance caused by lack of albumin free thiol triggers NETosis in an inflammation-independent manner.

### Plasma redox imbalance triggers lung-predominant NETosis

Next, we assessed the distribution of NETs deposited in the organs of mice with plasma redox imbalance. As the brain, lung, liver, and bone are known organs prone to metastasis formation for many cancer types, we assessed these four organs for evidence of NETs deposition in our mouse models of albumin deficiency and oxidation (i.e., Alb^−/−^ mice and IAA-injected Alb^wt/wt^ mice). Two-photon microscopy of these organs demonstrated that NET-like DNA structures stained by the cell-impermeable DNA dye, SytoxGreen, were detected exclusively in the lungs from Alb^−/−^ mice and from Alb^wt/wt^ mice 2 days post IAA injection (Fig. 3a, Extended Data Fig. 6a). These NET-like structures were not observed in the lungs of mice treated with DNase I, Cl-amidine, or albumin (Fig. 3b, Extended Data Fig. 6b), indicating that these SytoxGreen-positive web-like structures were compatible with NETs. We did not observe NETs within the other organs, including the liver, which instead only displayed round-shaped structures with green autofluorescence (Fig. 3a, Extended Data Fig. 6a) even in the absence of SytoxGreen (Extended Data Fig. 6c), compatible with vitamin A deposits^23^. In the extracts of the organs, DNA-associated Cit-H3 was enriched in those of the lung from Alb^−/−^ mice and IAA-injected Alb^wt/wt^ mice (Fig. 3c). Collectively, plasma redox imbalance induced by loss of albumin free thiol causes lung-predominant NETosis.

**Figure 3.**
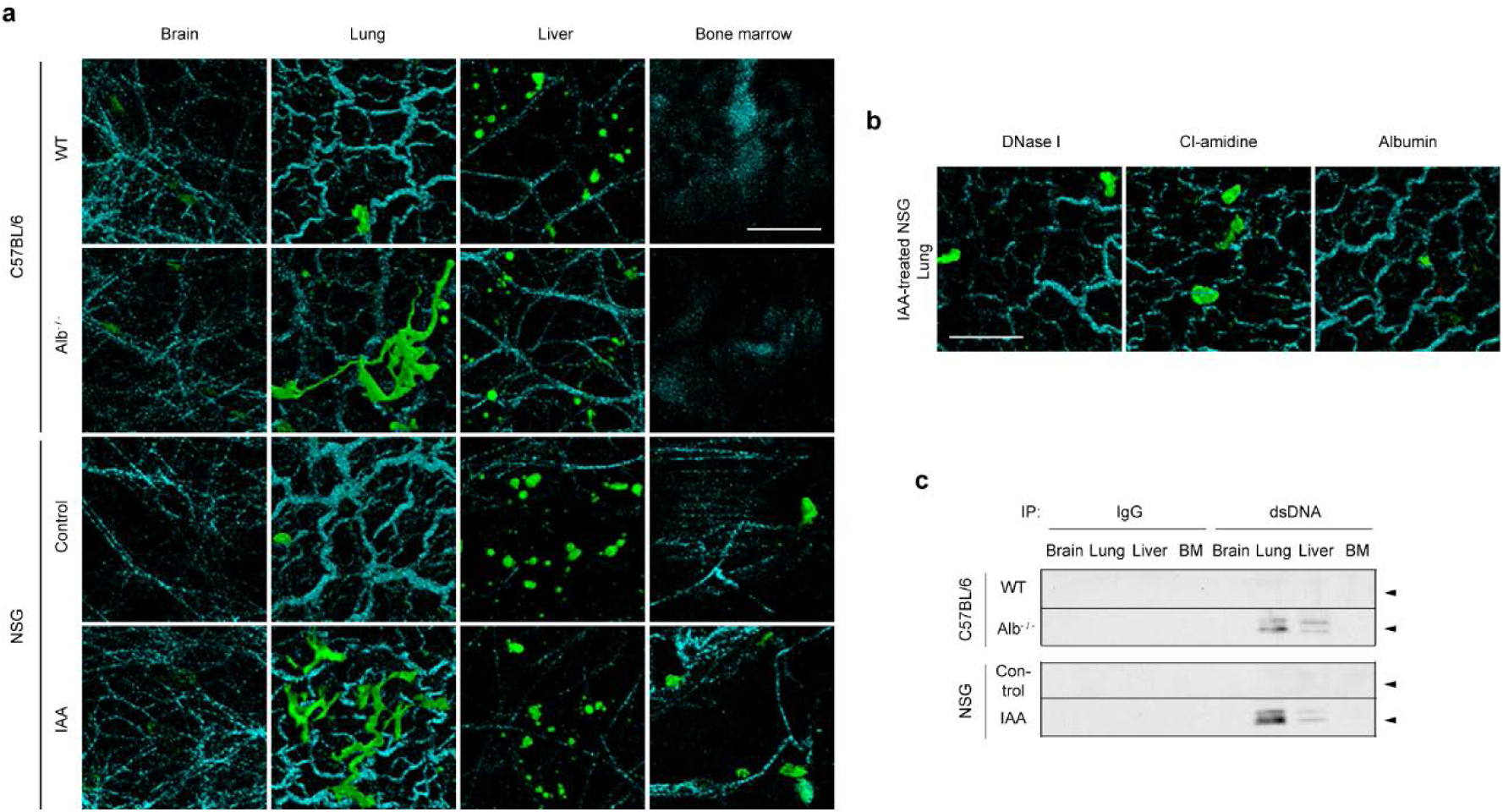
NETosis triggered by albumin deficiency or oxidation is lung-predominant. (a-c) The indicated organs were resected from WT and Alb^−/−^ C57BL/6 mice and NSG mice 2 days post saline or IAA (30 mg/kg) injection with or without single injection of DNase I on day 2, daily injection of Cl-amidine, or single injection of murine albumin on day 0. (a, b) The resected organs were subjected to two-photon microscopy. Extracellular DNA (green) and surrounding collagen-rich parenchyma (cyan) were visualized by SytoxGreen injected *via* tail vein 20 minutes before organ harvest and by second-harmonic generation, respectively. (a) Representative images are shown. Bar = 40 μm. (b) Representative images of the lungs from IAA-injected NSG mice with DNase I, Cl-amidine, or albumin treatment. Bar = 40 μm. (c) Extracts of the organs were subjected to immunoprecipitation using an anti-dsDNA antibody followed by Western blotting using an antibody against Cit-H3. Arrows indicate the molecular weight corresponding to Cit-H3.

### Plasma redox imbalance leads to elevated risk of pulmonary cancer metastasis via NETs

We next examined whether lung-predominant NETs generated by plasma redox imbalance promote pulmonary cancer metastasis through NET-dependent entrapping of CTCs. For this purpose, we used the mouse model of albumin oxidation by IAA rather than the Alb^−/−^ mouse because albumin has a wide variety of physiological functions (e.g., maintenance of intravascular pressure, drug transport, etc.)^24^ and its deficiency could affect cancer progression in various aspects. CAL-33 head and neck squamous cell carcinoma (HNSCC) cells with stable expression of either mCherry or *firefly* luciferase (CAL-33-mCherry and CAL-33-Luc, respectively) were intravenously injected into the tail vein of mice with or without IAA administration (Fig. 4a). Two-photon microscopy of *ex vivo* lungs harvested just after the injection of CAL-33-mCherry demonstrated greater numbers of trapped CTCs in IAA-injected *vs*. mock-injected mice (Fig. 4b, c). Clusters of CAL-33-mCherry cells were observed in close proximity to SytoxGreen-stained NETs in IAA-injected mice (Fig. 4b), but not in mice treated with DNase I or Cl-amidine to inhibit NET formation (Fig. 4b, c). Longitudinal observation of the trapped CTCs by optical imaging after the intravenous injection of CAL-33-Luc showed that mice with IAA-induced NETs displayed consistently elevated bioluminescent intensity in the lungs, while both DNase I and Cl-amidine reversed the effect of IAA (Fig. 4d, e). Of note, this mouse model of lung metastasis also showed sacrum bone metastasis (Extended Data Fig. 7a), but unlike the lung metastases NET induction did not have an effect on bone metastases (Extended Data Fig. 7b, c). These results indicate that lung-predominant deposition of NETs following IAA administration specifically promotes pulmonary metastases.

**Figure 4.**
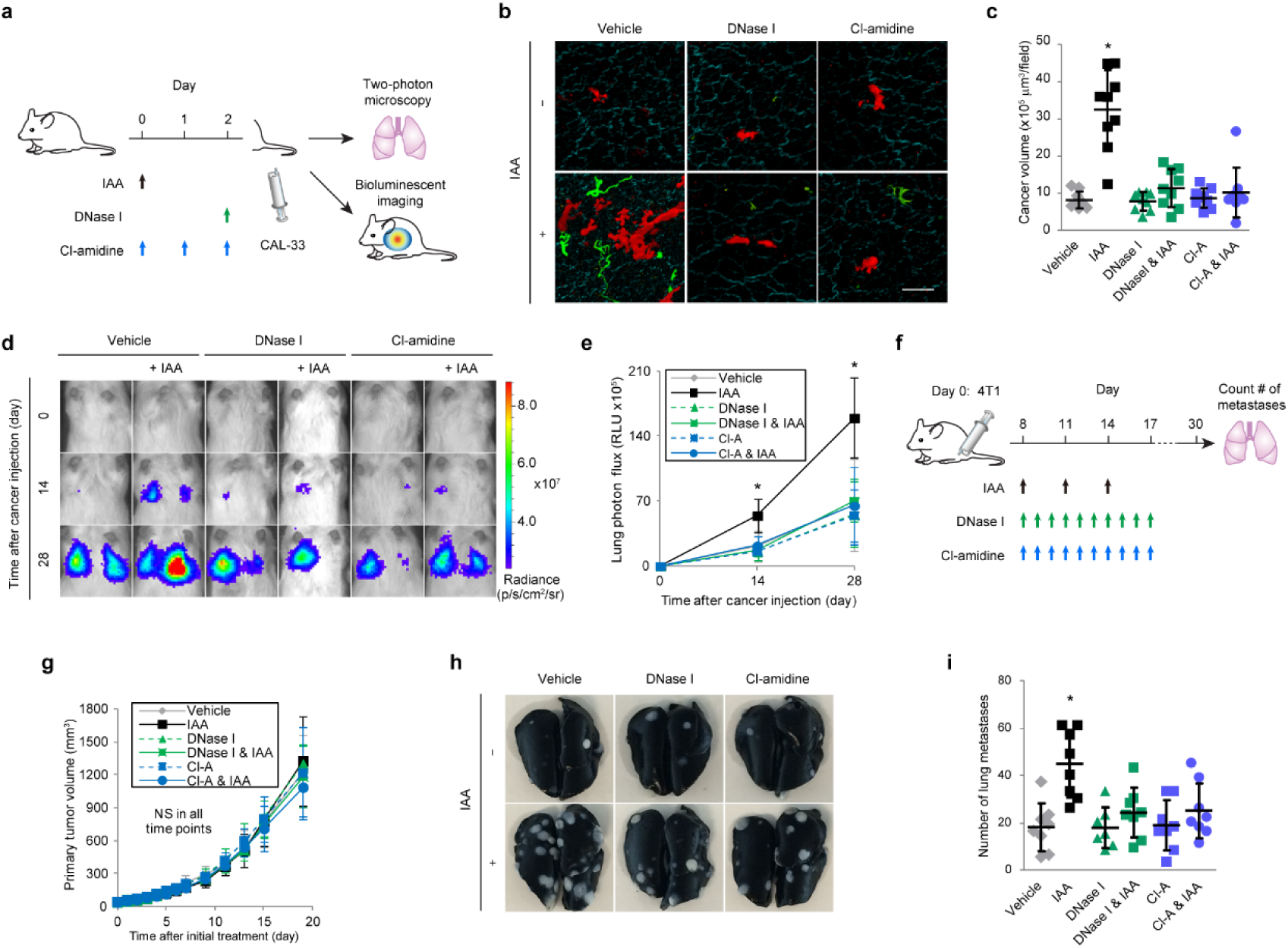
Plasma redox imbalance induced by albumin oxidation promotes lung metastasis *via* NETs. (a) Experimental schema for (b-e): the effect of NETs induced by albumin-thiol blocking on pulmonary metastasis following tail vein injection of CAL-33 HNSCC cells. Saline or IAA (30 mg/kg) was injected intraperitoneally into NSG mice. DNase I or Cl-amidine was administered at the indicated time points. Two day later, SytoxGreen and CAL-33-mCherry cells (b,c) and CAL-33-luciferase (d, e) were injected. (b, c) Extracellular DNA (green), CAL-33-mCherry cells (red), and surrounding collagen-rich parenchyma (cyan) within resected lungs were visualized by two-photon microscopy. (b) Representative images are shown. Bar = 50 μm. (c) Quantification of the mCherry-positive volume from two-photon microscopy images of resected lungs.(d, e) The growth of pulmonary metastases was monitored by bioluminescence imaging. (d) Representative images are shown. (e) Luciferase bioluminescence intensity from the lungs over time (0„28 days). (f) Experimental schema for (g-i): the effect of NETs induced by albumin-thiol blocking on a spontaneous pulmonary metastasis model. 4T1 murine mammary carcinoma cells were orthotopically transplanted to the mammary fat pads of female Balb/c mice. Saline or IAA (30 mg/kg) was injected intraperitoneally at the indicated time points. Between 8 to 17 days after transplantation, saline, DNase I, or Cl-amidine was administered daily. (g) Primary tumor growth. NS = not significant. (h, i) Lungs were removed 30 days after transplantation. (h) Representative images are shown. (i) Number of metastatic lung colonies. (c, i) Results represent individual values with the mean ± s.d.. (e, g) Results represent the mean ± s.d.. (c) *n* = 9 per group: 3 fields in 3 independent experiments. (e) *n*= 5 per group; one of the mice in the DNase I & IAA group died on day 1 for unknown reasons. (g, i) *n* = 8 per group; one of the mice in the DNase I group died on day 14 for unknown reasons. \**P*<0.05 by Dunnett’s test.

To further confirm these findings, a murine model of spontaneous pulmonary metastasis was utilized, wherein murine 4T1 breast cancer cells were transplanted orthotopically into the mammary fat pad of immunocompetent BALB/cByJ mice (Fig. 4f). Although the administration of IAA did not promote the growth of the primary tumor in the mammary fat pad (Fig. 4g), the number of metastatic colonies was markedly increased by IAA administration (Fig. 4h, i). This effect was rescued by treatment with DNase I or Cl-amidine (Fig. 4h, i), indicating its dependence on NETs. No other organs were affected by cancer metastasis (data not shown). Thus, our results from two distinct pre-clinical models indicate that albumin oxidation-triggered NETs in the lungs promote colonization of CTCs and initiation of pulmonary metastasis.

### Plasma redox imbalance is associated with inflammation-independent NETosis and pulmonary metastasis in a cohort of head and neck cancer patients

To investigate the association among plasma redox balance, NETosis, and lung metastasis in human cancer, we analyzed pre- and mid-treatment plasma samples from a cohort of 22 non-metastatic HNSCC patients undergoing curative therapy (Supplementary Table S1). Patients who developed subsequent lung metastasis (*n* = 8; median time-to-metastasis = 489 days [range: 115-786 days]) had a significantly lower concentration of plasma free thiol at mid-treatment compared to the 14 group-matched controls who did not develop metastasis (Fig. 5a; *p* = 0.001). Consistent with this result, significantly lower concentration of non-oxidized albumin (i.e., albumin containing a free thiol residue) in plasma was observed amongst the patients who subsequently developed lung metastasis (Fig. 5b; *p* = 0.011). Across the entire cohort, the concentration of non-oxidized albumin was positively correlated with free thiol concentration (R^2^ = 0.23, *p* = 0.001; Fig. 5c). Stratified at the median, HNSCC patients with <0.43 mM of free thiol or <2.4 g/dL of non-oxidized albumin at mid-treatment demonstrated significantly inferior lung metastasis-free survival (hazard ratio = 5.3 [95% CI 1.1„26.7] and 9.8 [95% CI 1.2„80.0], respectively; Fig. 5d, e) compared to patients with higher than the median values. Furthermore, a significant increase in NETs was observed in the patients with <2.4 g/dL vs. ≥2.4 g/dL of non-oxidized albumin at mid-treatment (Fig. 5f; *p* = 0.008). Between these groups, there was no significant difference in plasma concentration of CRP (Fig. 5g). Thus, albumin oxidation-associated decrease in plasma free thiols (i.e., plasma redox imbalance) during treatment could be an independent risk factor for developing elevated levels of NETosis and subsequent lung metastasis.

**Figure 5.**
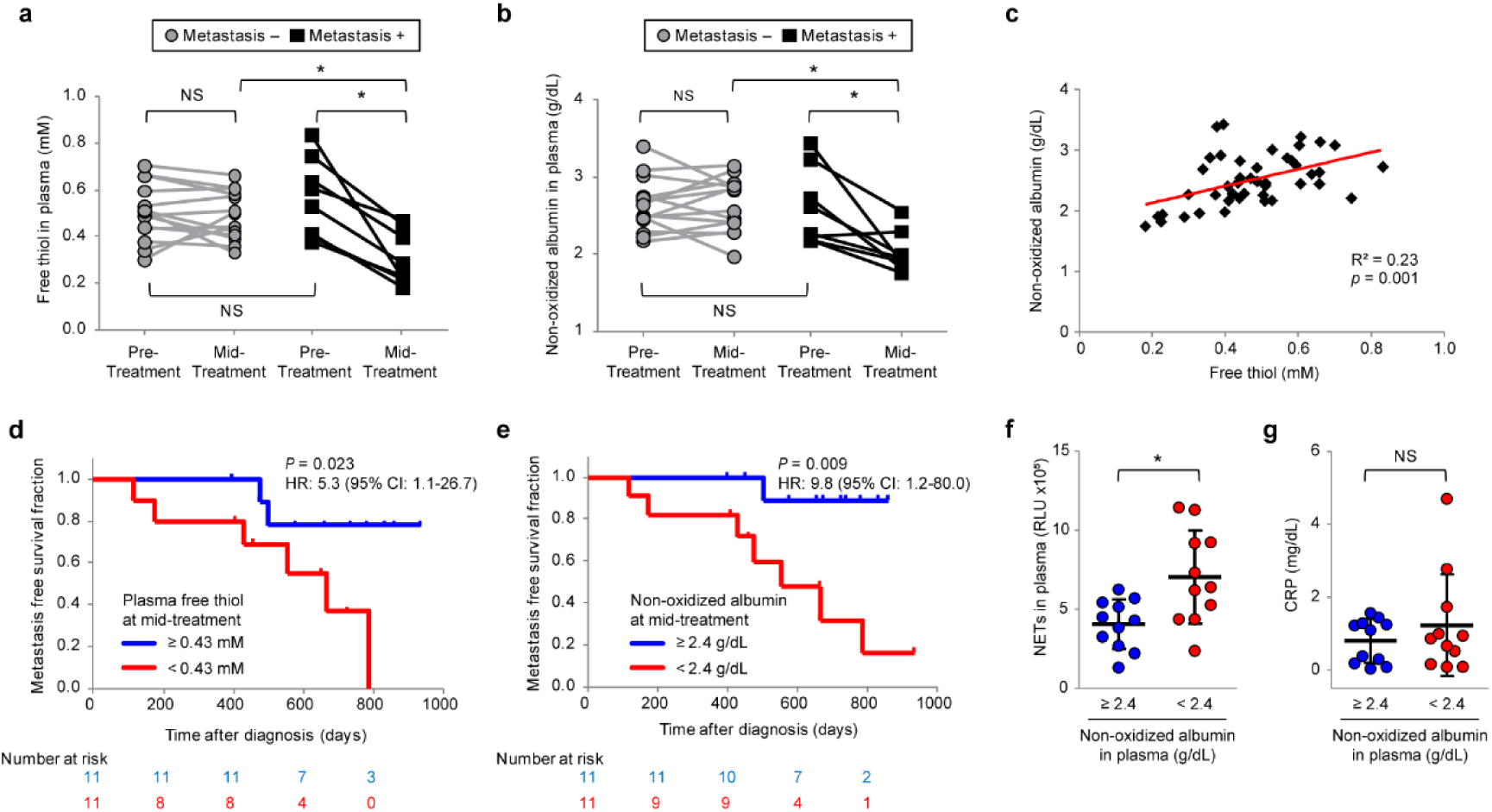
Plasma redox imbalance is associated with inflammation-independent NETosis and pulmonary metastasis in a human cancer cohort. (a-g) Pre- and mid-treatment plasma samples from HNSCC patients who developed lung metastasis (Metastasis +; *n* = 8) and matched control patients without metastasis (Metastasis „; *n* = 14) were analyzed for free thiol (a), and non-oxidized albumin (b) concentration. (c) Relationship between the concentration of plasma free thiol and non-oxidized albumin. The correlation between plasma free thiol and non-oxidized albumin was analyzed (*n* = 44, Pearson’s correlation coefficient test, R^2^ = 0.23, *p* = 0.001). (d, e) Kaplan-Meier analysis of lung metastasis-free survival in the entire cohort stratified by the median values of plasma free thiol (d), and non-oxidized albumin (e)concentration at mid-treatment (hazard ratio = 5.3 [95% CI 1.1„26.7] and 9.8 [95% CI 1.2„80.0], respectively; log-rank *p* = 0.023 and 0.009, respectively) (f, g) Plasma level of NETs (f) and CRP (g) in the HNSCC patient cohort stratified by the median values of non-oxidized albumin. The data are individual values with the mean ± s.d. (*n* = 11 in each group). \**P*<0.05 by (a, b) repeated-measures ANOVA for split-plot designs and (f, g) Student’s *t*-test, NS = not significant.

## Discussion

Until now, inflammatory-independent regulation of NETosis in human cancer has remained elusive. We have demonstrated that plasma redox imbalance caused by albumin oxidation triggers NETosis in an inflammation-independent manner. Our results further revealed a unique anatomical distribution of NET formation—predominantly in the lungs—that may further clarify the clinical significance of this phenomenon in cancer patients. Based on our findings, we propose that NET deposition within the lungs leads to entrapping of CTCs and ultimately pulmonary metastasis. Lung specificity could be explained by the fact that the pulmonary capillary bed is the largest vascular bed in the body and is the predominant site of physiological neutrophil margination^25^.

Oxidation of albumin thiols within the plasma of cancer patients may occur through multiple mechanisms. Known physiologic factors impacting plasma redox balance in cancer patients could include ROS produced by cancer cells^26^ and/or within the cancer microenvironment^27^. In addition, anti-cancer treatments such as radiotherapy and chemotherapy provide a repeated source of ROS generation^28^. We propose a model in which albumin, under normal physiologic conditions, provides an abundant source of free thiol within plasma that is available to scavenge ROS within circulating neutrophils. However, when this reservoir of albumin thiols becomes depleted in a cancer state, ROS may accumulate within neutrophils leading to NETosis and seeding of hematogenous metastases.

Our findings have significant implications for future clinical investigations. The relationship between neutrophils and cancer progression has been extensively studied^29^, but therapeutic targets that modulate neutrophil activity have not yet been identified. Herein, we propose that patients at elevated risk of distant metastasis might benefit from anti-NET therapies, such as DNase I or PAD inhibitors. This strategy has the potential for broad applications in oncology since, as our results demonstrate, even cancer patients under inflammation-free conditions may still be prone to develop NETs. Levels of plasma free thiol and/or non-oxidized albumin may provide a potentially generalizable biomarker that can assess the efficacy of such drugs in reversing the detrimental effects of dysregulated redox balance. Moreover, plasma redox balance may represent a novel and promising therapeutic target for NET inhibition and metastasis prevention. Together, the anti-NET therapies and potential predictive biomarkers described here could provide a potent clinical strategy to combat cancer metastasis.

## Methods

### Neutrophil isolation

Human whole blood from healthy volunteers was collected into EDTA-containing tube. Neutrophils were isolated by density gradient centrifugation using Polymorphprep (Alere Technologies, Germany) according to the manufacturer’s instructions. Neutrophils were re-suspended with Roswell Park Memorial Institute (RPMI) 1640 (Gibco/Life Technologies, MA, USA) without phenol red supplemented with 1% fetal bovine serum (FBS; Gibco/Life Technologies, MA, USA) for normal culture conditions. Alternatively, neutrophils were cultured under serum-free (i.e., RPMI-1640 with 0% FBS) or albumin-depleted (i.e., RPMI-1640 with 1% FBS without albumin – see *Albumin depletion from serum or plasma*) conditions. For the replenishment of albumin, bovine serum albumin (BSA; BioShop Canada Inc., Canada) dissolved in phosphate buffered saline (PBS) was freshly prepared and added to albumin-free media at a final concentration of 0.02 g/dL, matching the albumin concentration in the condition of 1% FBS. Neutrophil purity was established to be routinely >90%, as assessed by May-Grünwald Giemsa (Sigma-Aldrich, Canada) staining.

### Cell culture

The human head and neck squamous cell carcinoma (HNSCC) cell line CAL-33 was obtained from the American Type Culture Collection (VA, USA). The murine breast cancer cell line 4T1 and the embryonic human kidney cell line HEK293T were kind gifts from Dr. Graham Fletcher (Campbell Family Institute for Breast Cancer Research, Canada) and Dr. Bradly Wouters (Princess Margaret Cancer Centre, Canada), respectively. CAL-33 and HEK293T were cultured in 10% FBS-containing Dulbecco’s modified Eagle’s medium (Gibco/Life Technologies, MA, USA), and 4T1 cells were cultured in 10% FBS-containing RPMI-1640 in a well-humidified incubator with 5% CO_2_ and 95% air at 37°C.

### Albumin depletion from serum and plasma

Albumin was depleted from serum or plasma using polystyrene columns (ThermoFisher Scientific, MA, USA) packed with albumin-binding resin (CaptureSelect MultiSpecies Albumin Depletion; ThermoFisher Scientific, MA, USA). Five hundred microliters of serum or plasma was loaded into the column packed with 2.5 mL of resin and washed with PBS by gravity flow. Flow-through fractions were collected and immediately passed through a 0.22 μm sterile filter (MilliporeSigma, MA, USA) to remove bacterial contamination. The degree of albumin depletion was confirmed by quantifying the concentration of total protein and albumin with the Bradford protein assay (Bradford Reagent, Sigma-Aldrich, Canada) and bromocresol green assay (BCG albumin assay kit, Sigma-Aldrich, Canada), respectively. Adsorbed albumin was eluted with 0.1 M glycine, pH 3.0.

### Induction and inhibition of NETosis *in vitro*

For the induction of NETosis, human neutrophils (5×10^5^) cultured with RPMI-1640 containing 0% FBS, 1%FBS without albumin, or 1% FBS with 100 nM of phorbol 12-myristate 13-acetate (PMA; Cayman Chemical, MI, USA). For the inhibition of NETosis, human neutrophils (5×10^5^) were cultured with albumin-free media containing 0.02 g/dl of BSA, 1mM of *N*-acetylcysteine (NAC; Sigma-Aldrich, Canada), or 50 μM of Cl-amidine (MilliporeSigma, MA, USA). For NET inhibition by Cl-amidine, neutrophils in the media containing 1% FBS were pre-treated with 50 μM of Cl-amidine for 30 minutes. After six-hour incubation, the culture medium was collected after vigorous agitation. The medium was centrifuged at 100 x g for 1 min and the supernatant was used for the quantification of NETs.

### Quantification of extracellular and cell-free DNA

The concentrations of total extracellular DNA in culture medium and circulating cell-free DNA in plasma were determined by quantitative polymerase chain reaction (qPCR) with primers targeting the second open reading frame of the long interspersed element-1 transposon (LINE-1) (forward/reverse primer for human and mouse LINE-1: 5’-TCACTCAAAGCCGCTCAACTAC-3’ / 5’-TCTGCCTTCATTTCGTTATGTACC-3’ and 5’-AATGGAAAGCCAACATTCACGTG-3’ / 5’-CCTTCCTTGACCAAGGTATCATTG-3’, respectively; Eurofins Genomics, Belgium). The culture medium and plasma were diluted 1:100 with DNA suspension buffer (TEKnova, CA, USA). The reaction mixture for each qPCR consisted of 4 μL of a diluted sample, 1 μL of 2.5 μM LINE-1 primer mixture, and 5 μL of Sso Advanced Universal SYBR Green Supermix (Bio-Rad Laboratories, CA, USA). Real-time PCR amplification was performed with a pre-cycling heat activation of DNA polymerase at 98°C for 3 min followed by 40 cycles of denaturation at 98°C for 10 sec, and annealing and extension at 60°C for 30 sec using a CFX384 Touch Real-Time PCR Detection System (Bio-Rad Laboratories, CA, USA). Absolute quantification of DNA in each sample was determined by a standard curve with serial dilutions of human genomic DNA (Promega, WI, USA).

### Imaging of NETs *in vitro*

Human neutrophils cultured under the designated condition for 6 hours were fixed with 4% paraformaldehyde (PFA) for 10 min and permeabilized with PBS containing 0.2% Tween20 for 10 min. After washing with PBS three times and incubating for 20 min in the SuperBlock (PBS) Blocking Buffer (ThermoFisher Scientific, MA, USA), cells were treated with a rabbit anti-LL-37 antibody (1:1000; Osenses Pty Ltd., Australia) for 1 hour at room temperature. After washing with PBS containing 0.1% Tween20 three times, LL-37 was labeled with Alexa Fluor 488 anti-rabbit IgG secondary antibody (1:1000; Invitrogen Inc., CA, USA) for 1 hour at room temperature. After an additional 3 washes in PBS containing 0.1% Tween20, DNA was stained with DAPI (NucBlue Fixed Cell ReadyProbes Reagent, ThermoFisher Scientific, MA, USA). Slides were mounted in ProLong Gold antifade reagent (ThermoFisher Scientific, MA, USA). Images were acquired with a Zeiss LSM700 confocal microscope (Carl Zeiss, Germany). Separately, for staining of live/dead neutrophils and NETs, Hoechst33342 (2 drops/mL; NucBlue Live Cell ReadyProbes Reagent, ThermoFisher Scientific, MA, USA) and SytoxOrange (1:1000; Invitrogen Inc., CA, USA) were added into the culture medium 6 hours after initiation of treatment. Neutrophils were then observed with EVOS FL Cell Imaging System (Life technologies, CA, USA).

### Immunoprecipitation and Western blotting

Mouse polyclonal antibodies against double stranded DNA (Abcam, MA, USA) and IgG (MilliporeSigma, MA, USA) were conjugated to magnetic beads following the manufacturer’s protocol (Dynabeads Antibody Coupling Kit, ThermoFisher Scientific, MA, USA). The mixture of 100 μL of culture medium or mouse plasma and 25 μL of antibody-conjugated beads solution or 500 μL of extracts derived from 20mg of mouse organs (i.e., brain, lung, liver, and bone marrow from femur and 50 μL of antibody-conjugated beads solution were incubated with shaking for 1 hour at room temperature. The supernatant was removed by a magnet. The residual pellet was washed with PBS and re-suspended with 25 μL of 2x Laemmli Sample Buffer (Bio-Rad Laboratories, CA, USA) and boiled for 5 min at 95°C. Sample solution was separated from the magnetic beads and 20 μL was loaded onto a 4-20% SDS-PAGE gel (Bio-Rad Laboratories, CA, USA), transferred to nitrocellulose membranes (Bio-Rad Laboratories, CA, USA), then blocked in 2.5% skim milk for 1 h. After washing three times with Tris-buffered saline containing 0.1% Tween 20 (TBST) for 5 min each, the membranes were incubated with rabbit polyclonal anti-LL-37 antibody (1:1000, Osenses Pty Ltd., Australia) for 1 hour at room temperature, or rabbit polyclonal anti-histone H3 (citrulline R2 + R8 + R17) antibody (1:500, Abcam, MA, USA) overnight at 4°C. After washing three times with TBST for 5 min each, the membranes were incubated with horseradish peroxidase conjugated anti-rabbit IgG (1:2000; Cell Signaling Technology, MA, USA) for 1 hour at room temperature. Membranes were washed three times with TBST, developed with Clarity Western ECL Substrate (Bio-Rad Laboratories, CA, USA) and exposed to X-ray film (HyBlot CL Autoradiography Film, Denville Scientific Inc., MA, USA).

### Intracellular ROS detection

Intracellular ROS were measured by the DCFDA Cellular ROS detection Assay Kit (Abcam, MA, USA) according to the manufacturer’s protocol. Isolated human neutrophils were treated with 20 μM DCFDA for 30 min. After centrifugation (400 x g for 5 min), neutrophils were re-suspended with the designated medium (i.e., RPMI-1640 with 1% FBS, 0% FBS, and 1% albumin-depleted FBS with or without 0.02 g/dl of BSA/oxidized BSA or 1 mM of NAC) and seeded onto 96-well plates at 1 × 10^5^ cells per well. The ROS level represented by the fluorescence intensity of DCFDA was quantified using a fluorescence plate reader (TECAN Infinite M200Pro, Tecan Group Ltd., Switzerland).

### Imaging of intracellular ROS and albumin in neutrophils

Human neutrophils (5×10^5^) were cultured on glass coverslips in 24-well culture plates under the designated condition for 30 min. For visualization of the internalization of supplemented BSA by neutrophils, we added albumin-fluorescein isothiocyanate conjugate (Sigma-Aldrich, Canada) in 1:9 ratio to unlabeled BSA. This BSA mixture was added into the culture medium at final concentration of 0.02 g/dL. To stain intracellular ROS, CellROX Deep Red Reagent (5 μM; ThermoFisher Scientific, MA, USA) was added into the culture medium. After washing three times with PBS, neutrophils were fixed with 4% PFA for 20 min. After an additional washing three times with PBS, DNA was stained with DAPI (NucBlue Fixed Cell ReadyProbes Reagent, ThermoFisher Scientific, MA, USA). Slides were mounted in ProLong Gold antifade reagent (ThermoFisher Scientific, MA, USA). Images were acquired with a Zeiss LSM700 confocal microscope (Carl Zeiss, Germany).

### Lentiviral transfection

The packaging plasmid (psPAX2), the envelope plasmid (PMD2G), and pLenti-CMV-Puro-LUC were kind gifts from Dr. Bradly Wouters (Princess Margaret Cancer Centre). pCDH-CMV-mCherry-T2A-Puro was a kind gift from Kazuhiro Oka (plasmid # 72264; Addgene, MA, USA). For lentiviral production, HEK293T cells were co-transfected with 21 μg of psPAX2,10.5 μg of PMD2G, and 21 μg of pCDH-CMV-mCherry-T2A-Puro or pLenti-CMV-Puro-LUC using the Lipofection transfection reagent (Invitrogen Inc., CA, USA). High-titer viral solutions were prepared and used for transduction into CAL-33 cells. Antibiotics selection with puromycin (2.0 μg/ml, Gibco) was performed for 5 days.

### Blocking of free thiol in albumin *in vitro*

In order to oxidize BSA, PBS solution of BSA (0.1 g/dL) was incubated with 100 mM of chloramine-T (Sigma-Aldrich, Canada) or 375 mM of iodoacetamide (IAA; Sigma-Aldrich, Canada) for 1h at 37°C under oxygen-saturated conditions. In order to remove excess chloramine-T or IAA from the albumin solution, the mixture was centrifuged with Amicon Ultra 10K centrifugal filter device (MilliporeSigma, MA, USA) and re-suspended with 450 μL of PBS, repeated three times.

### Detection of free thiol in albumin solution and plasma

Culture medium (x1) or human/mouse plasma (x100 dilution with PBS (pH 8.0)) containing 0.1 mM 5,5’-Dithiobis(2-nitrobenzoic acid) (Sigma-Aldrich, Canada) was incubated at 27°C for 30 min. Absorbance at 412 nm was measured with a spectrophotometer. For calculation of free thiol concentration, molar extinction coefficient of 14.15 mM^−1^ cm^−1^ at 412 nm was used^30^.

### Quantification of plasma concentration of albumin

The mixture of 2.5 μl of plasma, 2.5 μl of water, and 200 μl of BCG albumin assay reagent (BCG albumin assay kit, Sigma-Aldrich, Canada) was incubated in a 96-well plate for 5 minutes at room temperature. The absorbance at 620 nm was measured using a plate reader (TECAN Infinite M200Pro, Tecan Group Ltd., Switzerland); absolute quantification was determined by a standard curve with serial dilutions of albumin.

### Albumin-deficient mouse model

Albumin-deficient strain (C57BL/6J-Alb^em8Mvw^/MvwJ) was purchased from Jackson Laboratory (ME, USA) as homozygote breeding pairs and maintained at the animal facility in Princess Margaret Cancer Centre. Wild type control C57BL/6 mice were obtained from the Princess Margaret Cancer Centre Small Animal Facility accredited by the Canadian Council on Animal Care. The plasma or organs from 5-8-week old male and female mice were used.

### Induction of NETosis *in vivo*

Saline solution of IAA (30 mg/kg of body weight) or lipopolysaccharide (8 mg/kg of body weight; Sigma-Aldrich, Canada) was injected intraperitoneally into 8-week old NOD scid gamma (NSG) mice, which were bred in-house at the Princess Margaret Cancer Centre Small Animal Facility accredited by the Canadian Council on Animal Care. Where indicated, murine albumin isolated from NSG mouse plasma (see *Albumin depletion from serum or plasma*) was injected into the tail vein (20 mg of murine albumin in 200 μL of PBS) prior to the saline/IAA injection. Blood was collected from the submandibular plexus (Extended Data Fig. 4a, b) or *via* cardiac puncture (Fig. 2b-g) at designated time points. Immediately after blood collection, plasma was isolated by centrifugation (2,500 x g for 10 min at 4°C). To eliminate cellular debris, an additional centrifugation was performed (16,100 x g for 10 min at 4°C); isolated plasma was stored at −80°C until analysis.

### Quantification of the oxidized/non-oxidized forms of albumin

Ten microliters of human and mouse plasma were diluted with 20 μL of 0.10 M sodium phosphate, 0.3 M NaCl (pH 6.87) and filtered through a Spin-X Centrifuge Tube Filters (VWR, PA, USA). Fast protein liquid chromatography (FPLC) analysis was performed using ÄKTA FPLC (GE Healthcare, UK). Albumin was separated on a Shodex Asahipak ES-502N column (Showa Denko Co., Ltd., Japan) at 35°C. Elution was performed by means of a linear gradient with increasing ethanol concentrations from 0% to 5% for human albumin and from 0% to 1% for mouse albumin in 0.05 mol/L sodium acetate and 0.40 mol/L sodium sulfate mixture (pH 4.85) at a flow rate of 1.0 mL/min. The obtained FPLC profiles for human albumin were subjected to Gaussian curve fitting by Igor Pro 7 (WaveMetrics, OR, USA). The fractions for oxidized and non-oxidized albumin were estimated by dividing the area of each fraction by the total area. Plasma concentration of each form of albumin was calculated by multiplying the fraction for each form of albumin by the plasma concentration of total albumin. As the fractions for reduced and oxidized murine albumin were not separated enough to be analyzed by Gaussian curve-fit algorithms, we utilized the ratio of peak intensity for reduced/oxidized albumin to assess the redox state of murine albumin.

### Lung and sacrum metastasis model by tail vein injection with or without NETosis and anti-NET therapies

As shown in Figure 4a, to induce NETosis, IAA was injected intraperitoneally into 6-8-week-old NSG male mice 2 days prior to the tail vein injection of CAL-33-luciferase (1 × 10^6^ cells in 200 μL PBS). For anti-NET therapies, a single intravenous injection of DNase I (3000 units; Sigma-Aldrich, Canada) was administered on day 2; 6-hours prior to injection of cancer cells. Cl-amidine (10 mg/kg body weight; MilliporeSigma, MA, USA) was injected intraperitoneally on days 0, 1 and 2.

### Microcomputed tomography of mice

A GE eXplore Locus Ultra microCT (GE Healthcare, GB) was used for data-acquisition in prone position under isoflurane inhalation (a 16-second Anatomical Scan Protocol (total of 680 images) at 80 kV, 50 mA, using a 0.15-mm Cu Filter, to achieve approximately 150 micron resolution.). STTARR’s GPU Reconstruction UI program (in-house software) was used to reconstruct CT scans in DICOM format. Images were analyzed by Siemens Inveon Research Workplace 4.0 (Siemens, PA, USA).

### Spontaneous lung metastasis model of orthotopic breast cancer with or without NETosis and anti-NET therapies

The suspension of 4T1 cells (1 ×10^5^ in 50 μL PBS) was injected into the right fifth mammary fat pad of 6-8-week-old Balb/c female mice (Jackson Laboratory, ME, USA). Eight days after tumor implantation, when tumor became palpable, mice were grouped by block randomization according their primary tumor size. Intraperitoneal injection of IAA (30 mg/kg body weight) or saline was initiated on day 8 and repeated on days 11 and 14 (Fig. 4f). As anti-NET therapies, daily intraperitoneal injection of Cl-amidine (10 mg/kg body weight) or daily intravenous injection of DNase I (1200 units) was performed until day 17. At the first day of IAA or saline injection, Cl-amidine was administered 6-hour prior to injection of IAA or saline. Primary tumor volume was calculated as 0.5 × length × width^2^. Thirty days later, mice were euthanized and lungs were removed. A 20-gauge needle was inserted into the trachea and the lung was gradually inflated using 1 ml of PBS solution with India ink (15% (v/v); Speedball, NC, USA). For de-staining, the lung was immersed in Fekete’s solution (100 ml 70% ethanol (Sigma-Aldrich Canada), 10 ml 4% formaldehyde (Sigma-Aldrich Canada) and 5 ml 100% glacial acetic acid (Fisher Scientific, PA, USA). The metastatic nodules were counted by visual observation.

### Optical imaging of luminescence in lung and sacrum metastases *in vivo*

Optical *in vivo* imaging was performed using the Xenogen IVIS Spectrum (Caliper Life Sciences Inc., MA, USA). Bioluminescent images were acquired 9 min after intraperitoneal injection of luciferin (150mg/kg body weight, XenoLight D-Luciferin - K^+^ Salt Bioluminescent Substrate, PerkinElmer, MA, USA). During imaging, mice were anesthetized with 2% isoflurane gas in the oxygen flow (1.5 L/min). Signal intensities were quantified and analyzed using Living Image Software version 4.5 (PerkinElmer, MA, USA).

### Optical imaging of NETs in the mouse organs with two photon microscopy

PBS solution of SytoxGreen (5 μL in 200 μL PBS; Invitrogen Inc., CA, USA) was injected into the tail vein of WT and Alb^−/−^ C57BL/6 mice or NSG mice 2 days post saline, IAA (30 mg/kg) injection, followed by 5 × 10^5^ CAL-33-mCherry cells in PBS 20 minutes later (Fig. 4b, c). Mice were then euthanized and organs (i.e., brain, lungs, liver, and femur) were immediately resected. Femur was cut in half along the long axis and bone marrow was exposed. Two-photon imaging was conducted using a Zeiss LSM710 (Carl Zeiss, Germany) coupled with a W Plan-Apochromat 20x/1.0 DIC water immersion objective (Carl Zeiss, Germany) and a Chameleon femtosecond laser (Coherent Inc., CA, USA). The excitation wavelength was set to 810 nm. SytoxGreen- and mCherry-emitted photons were collected using two emission filters and a beam splitter. For visualizing collagen structures, signals for second harmonic generation were recorded between 400-410 nm. To visualize three-dimensional structure of NETs, Z-stack images were obtained at 3.0 μm intervals for 7-10 Z-steps (Fig. 3a, b, 4b, Extended Data Fig. 6a-c) or 1.5 μm intervals for 20 Z-steps (Fig. 4c) and processed with the ZEN software (Carl Zeiss, Germany). Tumor-infiltrated volume in the lung was quantified as the mCherry-positive volume by Image J software (NIH, MD, USA).

### Human HNSCC patient plasma analysis

All studies involving HNSCC patient specimens were approved by the Research Ethics Board of University Health Network. Patients treated with definitive intensity-modulated radiotherapy or chemoradiotherapy between 2014 and 2015 were identified from a prospective Anthology of Clinical Outcomes^31^. Serial blood samples obtained prior to commencement of treatment, and at mid-treatment were analyzed from 8 patients who experienced distant relapse in the lung, and 14 control patients without relapse selected by propensity score matching for age, gender, clinical cancer stage, and treatment modality (i.e. radiotherapy or chemoradiotherapy).

### Quantification of NETs in human plasma

A rabbit polyclonal anti-neutrophil elastase antibody (1:500; Abcam, MA, USA) was coated onto 96-well microtiter plates (ThermoFisher Scientific, MA, USA) overnight at 4°C. After blocking in 1% BSA for 1 hour, the mixture of 50 μL of human plasma and 50 μL of PBS containing 0.3% BSA, 0.05% Tween 20, and 5 mM EDTA was loaded per well and incubated at room temperature for 1 hour. After washing three times with PBS containing 0.05% Tween 20 and 5 mM EDTA (PBST-EDTA), a horseradish peroxidase conjugated anti-DNA antibody (1:100; Zymo Research Corporation, CA, USA) was added to each well and incubated at room temperature for 1 hour. After washing three times with PBST-EDTA, the peroxidase substrate (Glo Substrate Reagent Pack (DY993), Bio-Techne Corporation/R&D Systems, MN, USA) was added. The intensity of chemiluminescence was measured using a plate reader (TECAN Infinite M200Pro, Tecan Group Ltd., Switzerland).

### Detection and quantification of C-reacting protein (CRP) in plasma

For detecting CRP in mouse plasma, 2 μL of plasma diluted with 16 μL of PBS and6 μL of 4x Laemmli Sample Buffer (Bio-Rad Laboratories, CA, USA) was sonicated for 15 minutes and boiled for 10 min at 95°C. This mixture was subjected to Western blotting (see *Immunopreciptation and Western blotting*) using anti-C reactive protein antibody (1:200, Abcam, MA, USA). Concentration of CRP in human plasma was measured by Human C Reactive Protein ELISA Kit (Abcam, MA, USA) according to manufacturer’s protocol. Samples were diluted 1:50,000 and measured in duplicates.

### Statistical analyses

Data are presented as mean and standard deviation (s.d.) or individual values with or without mean and s.d.. The significance of difference between two independent subjects and among multiple subjects was determined using the Student’s *t*-test and Dunnett’s test, respectively. Differences over time were repeated-measures ANOVA for split-plot designs. Correlation was analyzed using the Pearson correlation coefficient test. The Kaplan„Meier method was used to evaluate distant-metastasis-free survival, and log-rank tests compared survival proportions among groups. Data were obtained from at least three independent experiments with 1-3 technical replicates. Western blot was performed twice. For the lung metastasis mouse model shown in Fig. 4g, h, and i, mice were grouped by block randomization according their primary tumor size before the initiation of the treatment. Other experiments were not randomized and there was no blinding. All tests were two-tailed with a *P* value <0.05 considered to be significant. All calculations and analyses were performed in R (version 2.8.1; http://www.R-project.org).

### Ethics of animal experiments

All animal experiments were approved by the Animal Research Committee of Princess Margaret Cancer Centre, and performed according to guidelines governing animal care in Canada.

## Acknowledgments

We appreciate the technical support of Reza Kiarash with the 4T1 experiments. This study was supported by a fellowship award to Min.I. from the EIRR21 program and Nakatomi foundation. S.V.B. is supported by the Gattuso-Slaight Personalized Cancer Medicine Fund at the Princess Margaret Cancer Centre. We also gratefully acknowledge the support from the Princess Margaret Cancer Foundation and the Princess Margaret Cancer Center Head & Neck Translational Program, with philanthropic funds from the Wharton Family, Joe’s Team, and Gordon Tozer.

## Author contributions

Min.I., M.E., Y.K., and S.V.B. conceived and designed the experiments; Min.I. performed the experiments; Min.I. and M.E. analyzed the data; Mit.I. contributed reagents and materials; Min.I. and S.V.B wrote the manuscript. All authors provided approval of the final manuscript.

## Additional information

Competing financial interests: The authors declare no competing financial interests.

---

**Supplementary Table S1.**
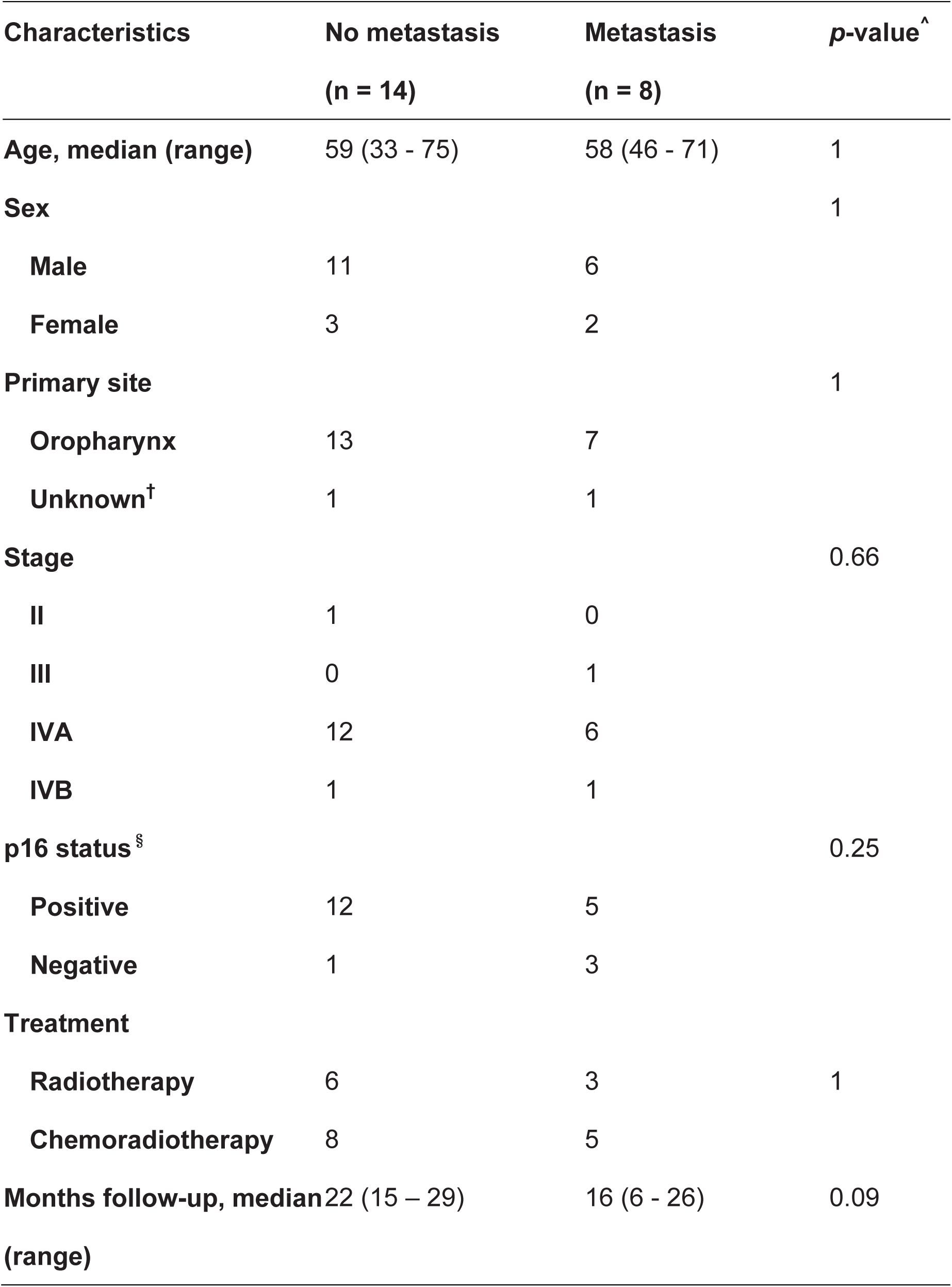
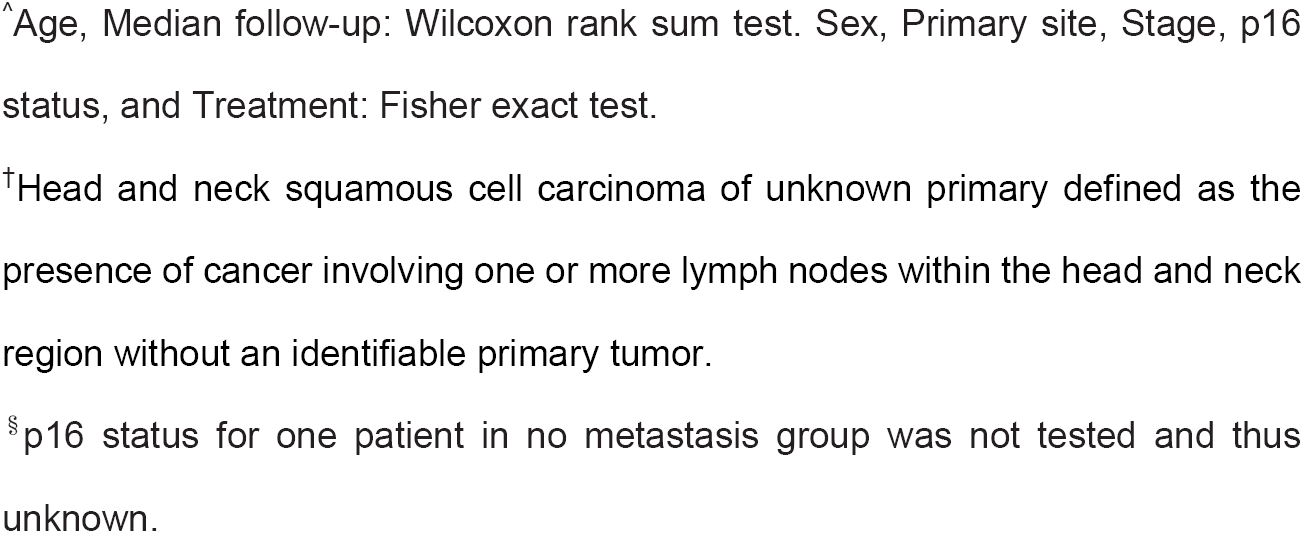
Patient characteristics.

| Characteristics | No metastasis<br>(n = 14) | Metastasis<br>(n = 8) | p-value<sup>^</sup> |
| --- | --- | --- | --- |
| **Age, median (range)** | 59 (33 - 75) | 58 (46 - 71) | 1 |
| **Sex** |  |  | 1 |
| Male | 11 | 6 |  |
| Female | 3 | 2 |  |
| **Primary site** |  |  | 1 |
| Oropharynx | 13 | 7 |  |
| Unknown<sup>†</sup> | 1 | 1 |  |
| **Stage** |  |  | 0.66 |
| II | 1 | 0 |  |
| III | 0 | 1 |  |
| IVA | 12 | 6 |  |
| IVB | 1 | 1 |  |
| **p16 status<sup>§</sup>** |  |  | 0.25 |
| Positive | 12 | 5 |  |
| Negative | 1 | 3 |  |
| **Treatment** |  |  |  |
| Radiotherapy | 6 | 3 | 1 |
| Chemoradiotherapy | 8 | 5 |  |
| **Months follow-up, median**<br>**(range)** | 22 (15 – 29) | 16 (6 - 26) | 0.09 |

<sup>^</sup>Age, Median follow-up: Wilcoxon rank sum test. Sex, Primary site, Stage, p16
status, and Treatment: Fisher exact test.
<sup>†</sup>Head and neck squamous cell carcinoma of unknown primary defined as the presence of cancer involving one or more lymph nodes within the head and neck region without an identifiable primary tumor.
<sup>§</sup>p16 status for one patient in no metastasis group was not tested and thus
unknown.

**Extended Data Figure 1.**
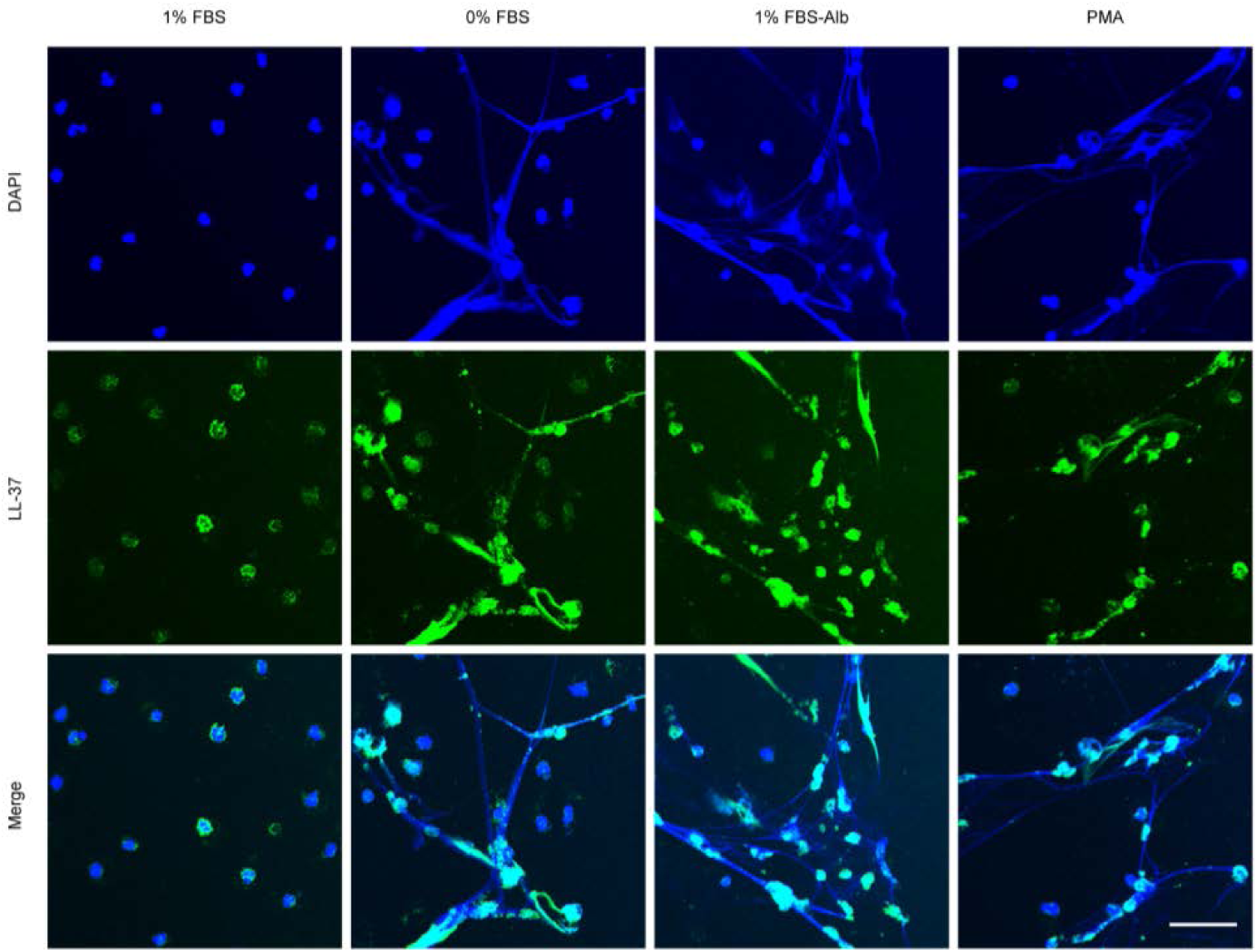
Immunocytochemistry for human neutrophils with or without albumin depletion. Human neutrophils were cultured in medium containing 1% FBS, 0% FBS, and 1% albumin-depleted FBS (1% FBS-Alb) for 6 hours. As a positive control for neutrophils undergoing NETosis, human neutrophils cultured in medium containing 1% FBS were treated with 100 nM of phorbol 12-myristate 13-acetate (PMA) for 6 hours. Representative immunofluorescence images are shown of neutrophils stained with DAPI (blue) and anti-LL37 (green). Bar = 50 μm.

**Extended Data Figure 2.**
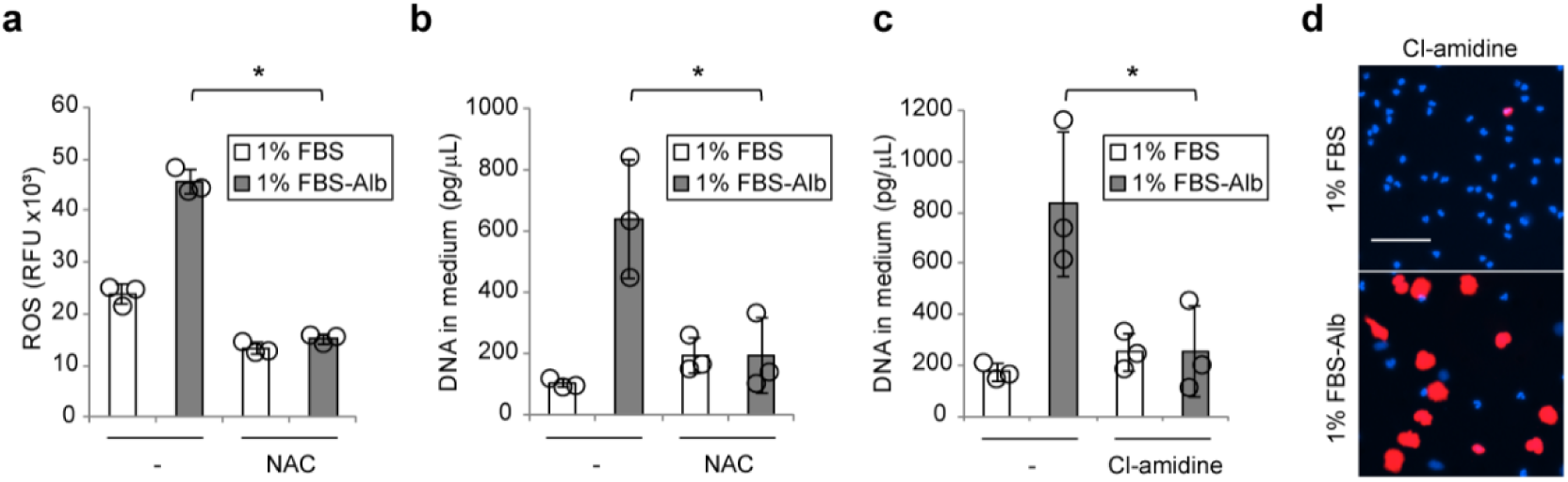
PAD inhibition and NAC supplementation prevent albumin depletion-evoked NETosis. (a-d) Human neutrophils were cultured in medium containing 1% FBS or 1% albumin-depleted FBS (1% FBS-Alb) with or without NAC (a, b) or a PAD inhibitor, Cl-amidine (c, d) or for 5 minutes (a) and 6 hours (b-d). (a) Intracellular ROS level. (b) Concentration of extracellular DNA within culture medium. (c) Concentration of extracellular DNA within culture medium. (d) Representative images of neutrophils stained with cell-permeable DNA dye, Hoechst 33342 (blue), and cell-impermeable DNA dye, SytoxOrange (red). Bar = 50 μm. (a-c) Results represent individual values with the mean ± s.d. (*n* = 3; biological triplicates. \**P*<0.05 by Student’s *t*-test). PAD: peptidyl arginine deiminase. NAC: *N*-acetylcysteine.

**Extended Data Figure 3.**
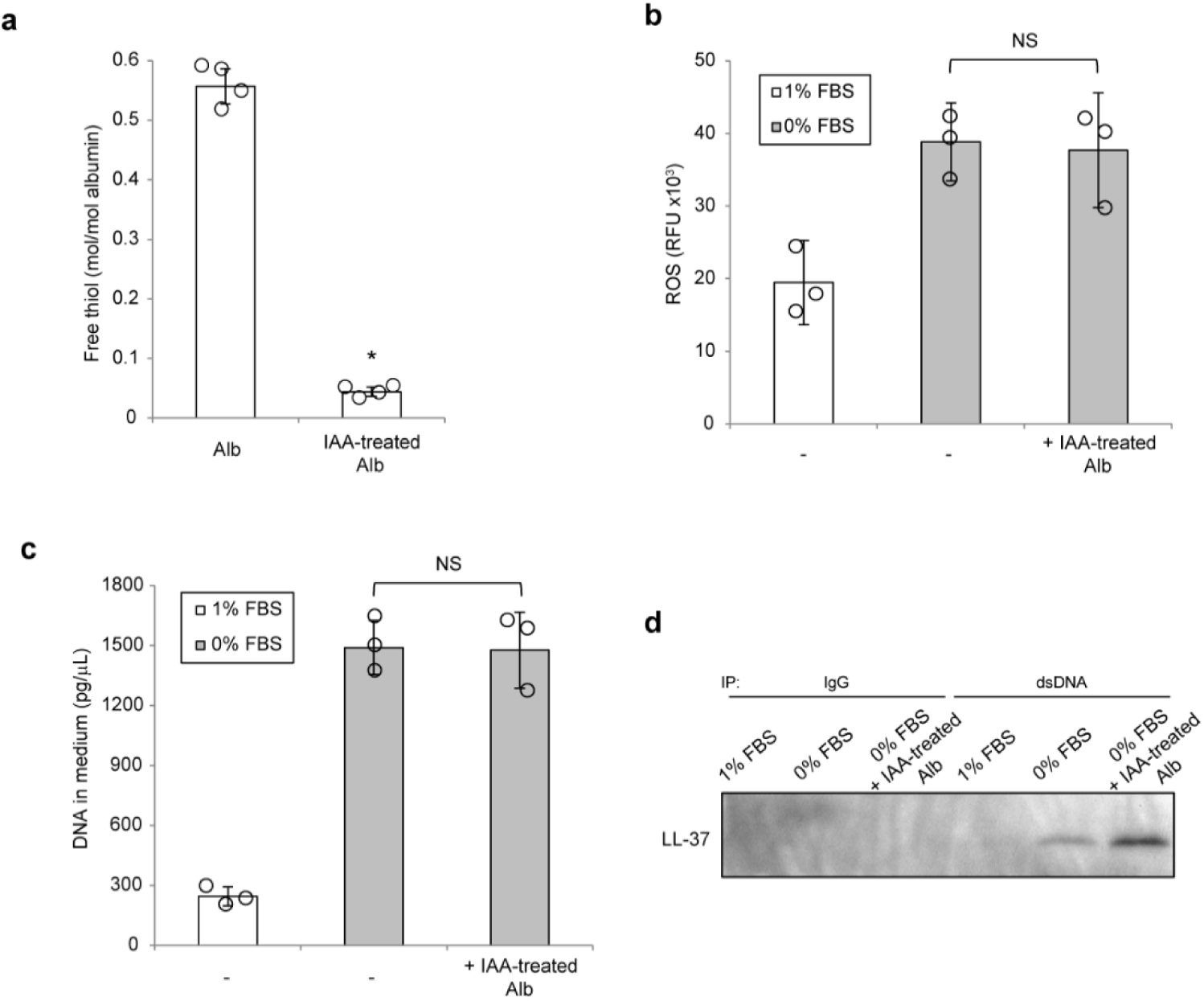
Iodoacetamide-treated albumin triggers NETosis through the accumulation of intracellular ROS. (a) Free thiol concentration in the BSA solution with or without free thiol blocking by IAA. (b-d) Neutrophils were 885 cultured in the indicated conditions. (b) Intracellular ROS level was quantified. (c) Concentration of extracellular DNA within culture medium. (d) Immunoprecipitated extracellular DNA was subjected to Western blotting for LL-37. (a-c) Results represent individual values with the mean ± s.d.. *n* = 3-4. \**P*<0.05. NS = not significant by Student’s *t*-test. BSA: bovine serum albumin.ROS: reactive oxygen species. IAA: iodoacetamide.

**Extended Data Figure 4.**
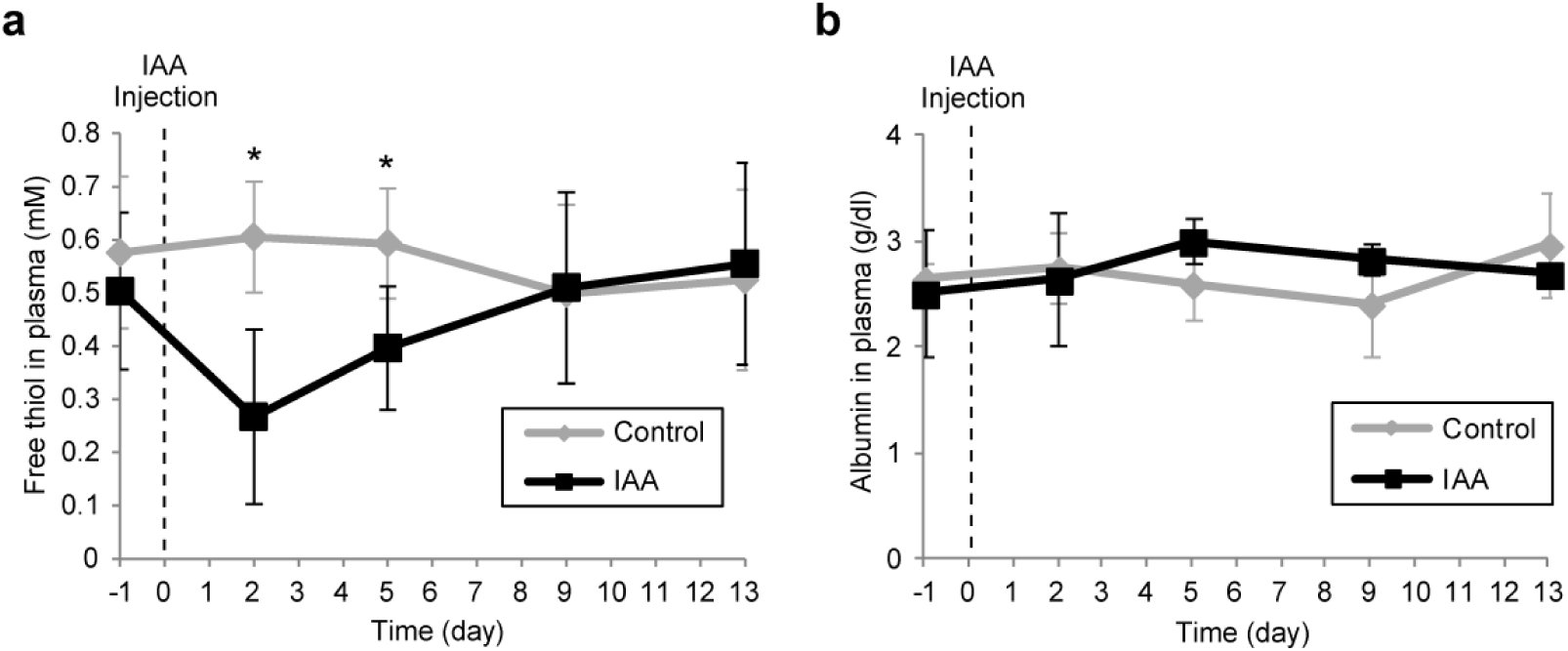
Determination of the optimal time point for the detection of NETosis by iodoacetamide. Saline or IAA was injected 895 intraperitoneally into NSG mice. Blood was collected at designated time point. Concentration of plasma free thiol (a) and albumin (b). (a-b) Results represent the mean ± s.d.. *n* = 3. \**P*<0.05 by Student’s *t*-test at each time point. IAA: iodoacetamide.

**Extended Data Figure 5.**
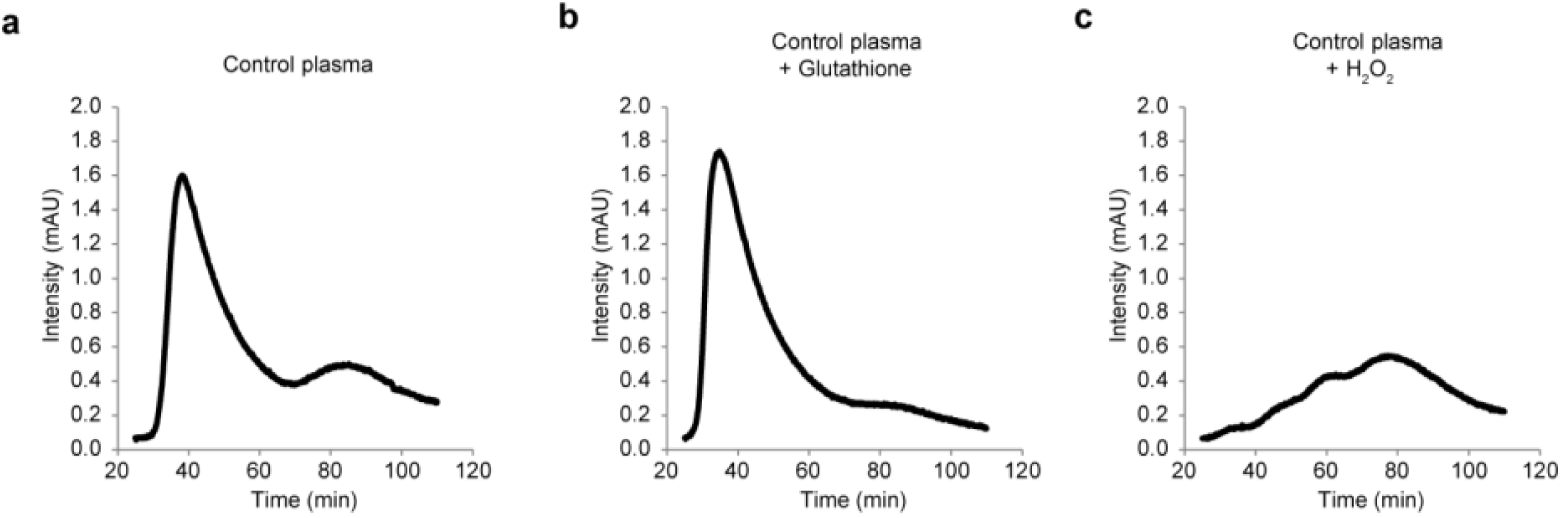
Liquid chromatography profile of murine albumin with exogenous reduction and oxidation. Murine albumin in plasma were analyzed by fast protein liquid chromatography. (a) Control mouse plasma. (b and c) Mouse plasma was incubated with 5 mM glutathione for 3 hours (b) or 50 mM hydrogen peroxide for 24 hours (c) to demonstrate that the first and second peaks represent the fraction of reduced and oxidized albumin, respectively.

**Extended Data Figure 6.**
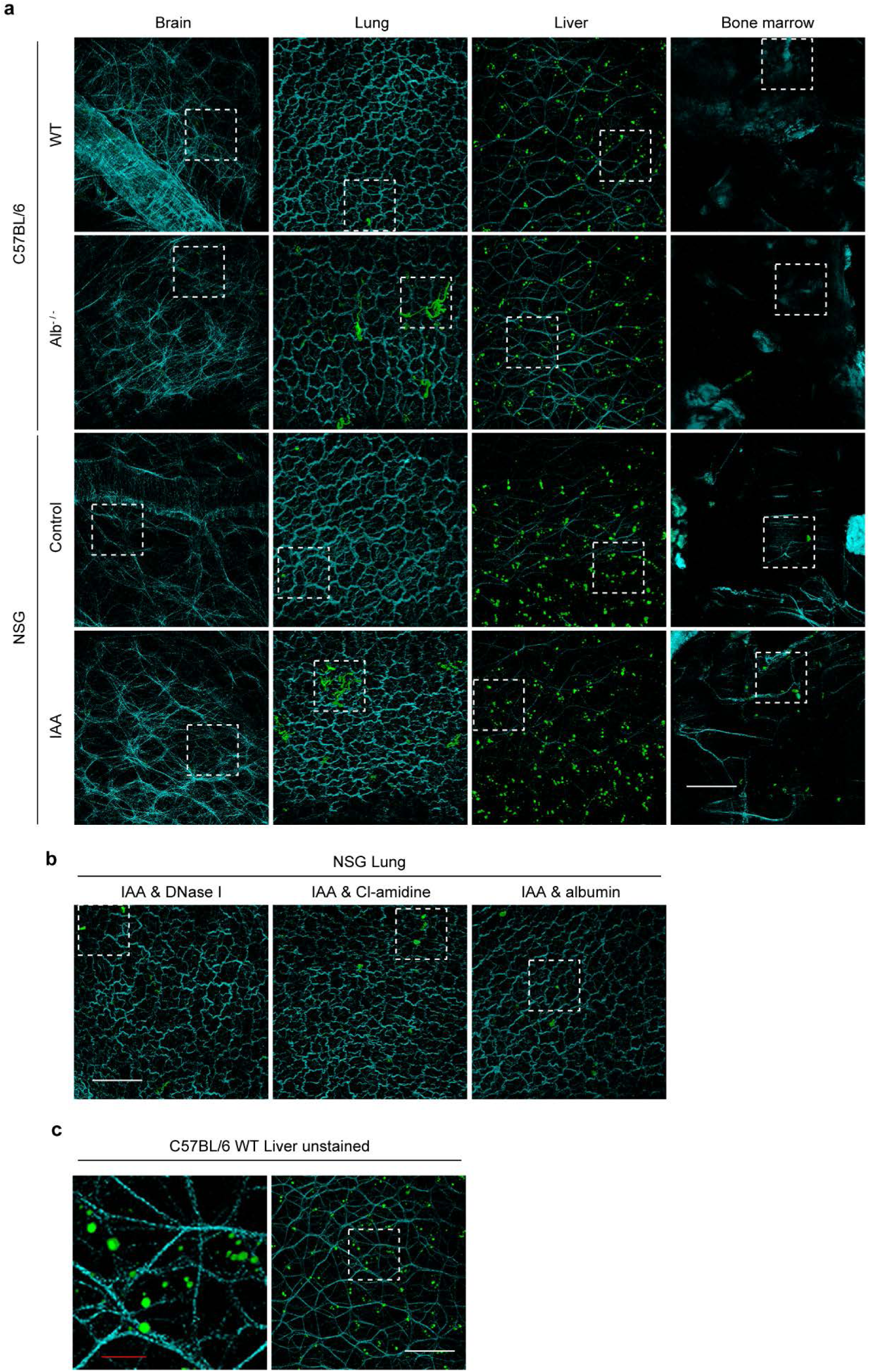
Albumin deficiency and albumin thiol-blockade induces lung-predominant NETosis. The indicated organs resected from wild-type (WT) and albumin deficient (Alb^−/−^) C57BL/6 mice and vehicle- and IAA-injected NSG mice were subjected to two-photon microscopy. Surrounding collagen-rich tissues were imaged by second-harmonic generation (shown as cyan). (a, b) To stain extracellular DNA, SytoxGreen was injected *via* tail vein 20 minutes before organ harvest (shown as green). (a) Representative images of brain, lung, liver, and bone marrow. The view in the dotted square is enlarged in Fig. 2h. Bar = 100 μm. (b) In order to inhibit NET formation, single injection of DNase I on day 2, daily injection of Cl-amidine, single injection of murine albumin on day 0 was performed in IAA-injected mice. Representative images of the lung were shown. The view in the dotted square is enlarged in Fig. 2j. Bar =100 μm (c) Representative image of the unstained liver from WT C57BL/6 untreated mice. The view in the dotted square in the right picture is enlarged in the left. Autofluorescence was shown as green. Red bar = 20 μm; White bar = 100 μm.

**Extended Data Figure 7.**
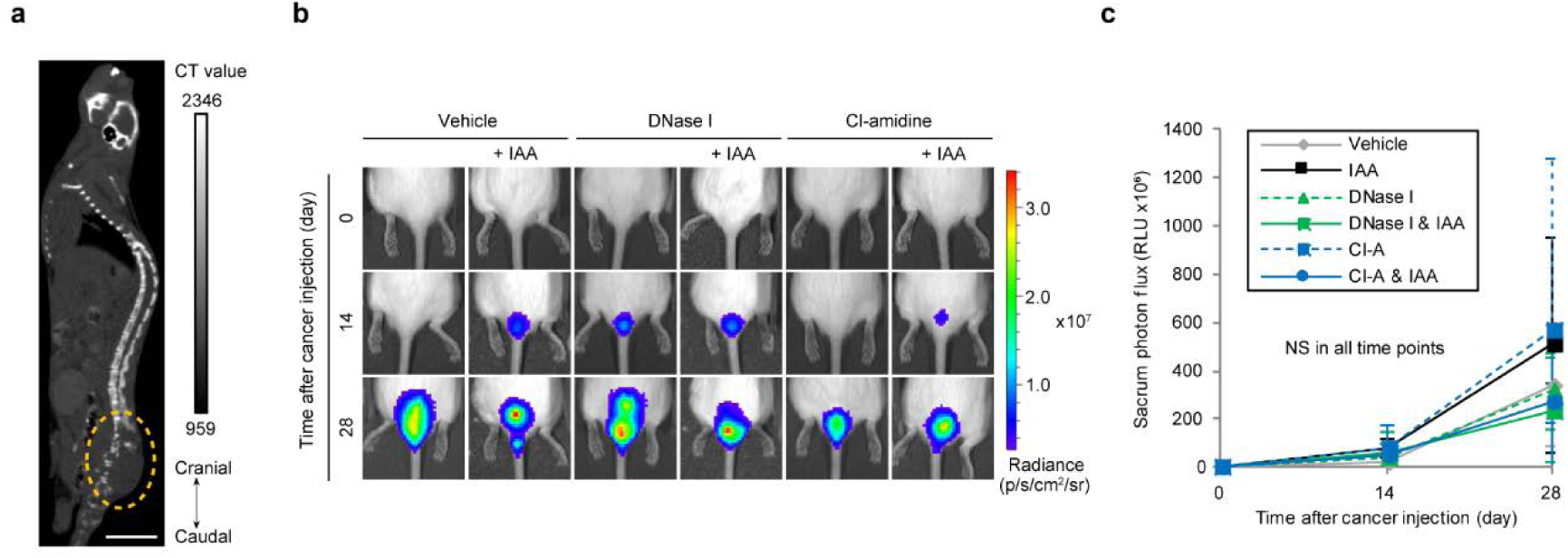
NETosis induced by albumin thiol-blockade does not promote bone metastasis. Saline or IAA was injected intraperitoneally into NSG mice. Two day later, CAL-33-luciferase were injected. The growth of sacrum metastases was monitored by bioluminescence imaging. (a) Representative image of computed tomography (CT) imaged 30 days after CAL-33-luciferase injection. The dotted ellipse represents the region of osteolytic sacrum metastasis. Bar = 1.5 cm. (b) Representative bioluminescent images are shown. (c) Luciferase bioluminescence intensity from the sacrum over time (0„28 days). *n* = 5 per group; one of the mice in the DNase I & IAA group died on day 1 for unknown reasons and was excluded. NS = not significant by Dunnett’s test.

